# Advancing Computational Biology and Bioinformatics Research Through Open Innovation Competitions

**DOI:** 10.1101/565481

**Authors:** Andrea Blasco, Michael G. Endres, Rinat A. Sergeev, Anup Jonchhe, Max Macaluso, Rajiv Narayan, Ted Natoli, Jin H. Paik, Bryan Briney, Chunlei Wu, Andrew I. Su, Aravind Subramanian, Karim R. Lakhani

**Affiliations:** Laboratory for Innovation Science at Harvard, Harvard University, Boston, MA 02163, USA; Harvard Business School, Harvard University, Boston, MA 02163, USA; Institute for Quantitative Social Science, Harvard University, Cambridge, MA 02138, USA; Broad Institute, Cambridge, MA 02142, USA; Department of Integrative Structural and Computational Biology, The Scripps Research Institute, La Jolla, CA 92037, USA; Department of Immunology and Microbial Science, The Scripps Research Institute, La Jolla, CA 92037, USA; National Bureau of Economic Research, Cambridge, MA 02138, USA

## Abstract

Open data science and algorithm development competitions offer a unique avenue for rapid discovery of better computational strategies. We highlight three examples in computational biology and bioinformatics research where the use of competitions has yielded significant performance gains over established algorithms. These include algorithms for antibody clustering, imputing gene expression data, and querying the Connectivity Map (CMap). Performance gains are evaluated quantitatively using realistic, albeit sanitized, data sets. The solutions produced through these competitions are then examined with respect to their utility and the prospects for implementation in the field. We present the decision process and competition design considerations that lead to these successful outcomes as a model for researchers who want to use competitions and non-domain crowds as collaborators to further their research.

## Introduction

Crowdsourcing enables large communities of individuals to collectively address a common challenge or problem. Mechanisms for crowdsourcing include tools to encourage voluntary work in data collection and curation, gamification of scientific problems, and the use of open innovation competitions. Researchers in computational biology increasingly rely on the first two mechanisms to tackle simple yet labor-intensive tasks, such as annotating images [1] or predicting aspects of protein structure [2]. Additionally, researchers use open innovation competitions to benchmark their solutions to a particular computational problem, such as the algorithm MegaBLAST for comparing DNA sequences [3], or to generalize their methodologies to instances of the problem for which a solution is unknown, such as inferring molecular networks [4].

Past examples demonstrate that open innovation competitions possess considerable potential in addressing biology problems [5]. However, most applications are intended for a “crowd” of researchers in communities that are most directly connected to the scientific problem at hand, such as the community of computational biologists to predict drug sensitivity [6]. Less is known about the potential of innovation competitions in biology when they are open to a crowd of *non-experts,* although there have been promising examples in other domains [7, 8]. This suggests a need for methodological developments on the use of competitions when the community of experts is small or nonexistent, or when it lacks the necessary knowledge to solve the problem (e.g., rapidly evolving or emerging fields that depend heavily on large amounts of data and computational resources but are deficient in experts in scientific computing, data science or machine learning).

In this study, we focus on contests that involve participants outside the community of researchers connected to the scientific problem, as in [3]. Rather than the prospect of advancing research in the field or achieving a scientific publication, researchers have to incentivize participation with opportunities to win cash prizes.^1^ Furthermore, they must articulate the problem in clear and easily-digestible language and construct metrics to evaluate solutions that provide live feedback to participants. Although competitors in these contests typically lack domain-specific expertise, they bring a breadth of knowledge across multiple disciplines, which creates the opportunity for exploration and cross-fertilization of ideas. Open innovation competitions of this nature allow members of the public to contribute meaningfully to academic fields by providing incentives and easy access to problems and data that are otherwise inaccessible to them.

We describe the design and outcomes of three contests in computational biology and bioinformatics to address problems whose solutions were significantly improved using open innovation competitions. These include algorithms for:

- clustering antibody sequences (Sec. 2.1);
- imputing gene expression measurements (Sec. 2.2);
- performing fast queries on the Connectivity Map dataset (Sec. 2.3).

The improvements achieved by these competitions focused on both optimization (reduction in speed or memory footprint) and production of better models (higher accuracy in statistical inference) and highlighted the versatility of crowdsourcing as an effective tool for researchers.

## 2 Results

The competitions discussed here were hosted on the platform Topcoder.com (Wipro, Bengaluru, India), providing access to a community of approximately one million software engineers and data scientists. They were held for a duration of two to four weeks, and they were structured so that each competitor could make multiple submissions and receive continuous feedback on the performance of their solution (based on objective metrics) summarized on a live public leaderboard. The purpose of such feedback was essential to drive competition among the participants. Incentives for these competitions consisted of cash prizes, ranging from $8,500 to $20,000. Specific payouts and participation levels for each competition are summarized in Table 1.

**Table 1:** Competition summary: Prize pools were distributed to the top-placing solutions, rank-ordered by performance based on objective evaluation metrics. For these contests, we used a constant prize pool over duration ratio (Prize Pool/Duration ~ 1000 USD/day) and the prize distributions follow: *P*(*N*) ~ *P*(1)*e*^-2/3(*N*-1)^ for places N = 1, 2,… and *P*(1) = Prize Pool/2.

| Contest | Duration<br>(days) | Prize Pool (\$1000)<br>(Distribution) | Participants | Average Submissions<br>per Participant |
| --- | --- | --- | --- | --- |
| Antibody Clustering | 10 | 8.5 (5/2/1/0.5) | 34 | 6.91 |
| Connectivity Map 1 | 21 | 20 (10/5/2.5/1.5/1) | 85 | 13.73 |
| Connectivity Map 2 | 21 | 20 (9/5/3/2/1) | 40 | 4.58 |

**Table 2:** Dataset descriptives for the CMap Inference Challenge.

| Dataset | Size (genes × samples) | Platform | Source |
| --- | --- | --- | --- |
| Current | $12,320 \times 12,000$ | Affymetrix | GEO |
| Training | $12,320 \times 100,000$ | Affymetrix | GEO |
| Testing | $970 \times 1,600$ | L1000 | GTE<sub>x</sub> |
| Ground | $12,320 \times 1,600$ | RNA-Seq | GTE<sub>x</sub> |

**Table 3:** Top five submissions for the CMap Inference Challenge.

| Rank | Algorithm description |
| --- | --- |
| 1 | KNN (average over multiple normalizations) |
| 2 | Neural Net with batch normalization |
| 3 | KNN (average over random subsets) |
| 5 | KNN (average over random subsets) |
| 5 | KNN (average over random subsets) |

We evaluated each submission using predetermined performance metrics (Online Methods, Sec. 6). These are real-valued functions of one or more characteristics, such as computation time, memory use, or accuracy, which targeted specific contest objectives. Three kinds of data sets were used for evaluation: a *training* set, a *validation* set, and a *test* set. The training set and associated ground truth were provided to participants for the purpose of understanding the properties of the data and for developing or training their solutions. The validation data set (excluding ground truth) was used to provide real-time feedback on the performance of their solution via submission of either source code or predictions produced by their algorithms. The test set was reserved for the end of the competition and was used for final evaluation of submissions; this data set was withheld from participants at all times. Note that for data science competitions, the holdout test set is particularly important for determining the true performance of solutions, since continuous feedback of performance in the form of a real-time leaderboard raises the prospect of model overfitting on the validation set.

### 2.1 Antibody Clustering Challenge (Scripps)

#### 2.1.1 Motivation and Objective

In the process of immunotherapy and vaccine development, next-generation sequencing of antibody repertoires allows researchers to profile the large pools of antibodies that comprise the immune memory formed following pathogen exposure or immunization. During the formation of immune memory, antibodies mature in an iterative process of clonal expansion, accumulation of somatic mutations, and selection of mutations that improve binding affinity. Thus, the memory response to a given pathogen or vaccine consists of “lineages” of antibodies which can trace each of their origins to a single precursor antibody sequence. Detailed study of such lineages can provide insight to the development of protective antibodies and the efficacy of vaccination strategies. After the initial alignment and assembly of reads, the antibody sequences can be clustered based on their similarity to the known gene segments encoding heavy or light antibody chains. Clustering these antibody sequences allows researchers to understand the lineage structure of all antibodies produced in individual B cells [9, 10, 11].

The number of antibody sequences from a single sample can easily reach to the millions; posing a major computational challenge for clustering at such a large scale. The bottleneck lies at computing a pairwise distance matrix and subsequently performing hierarchical clustering of the sequences. The former task scales as *O*(*N*^2^) in both computational and space complexity, where N is the total number of input sequences. The latter task, assuming a typical agglomerative hierarchical clustering algorithm, has a computational complexity that scales as *O*(*N*^2^ log *N*). By comparison, the file I/O is expected to scale as *O*(*N*).

#### 2.1.2 Benchmarks and Methodologies Prior to the Competition

Prior to the competition, a Python implementation of the clustering algorithm was developed, utilizing numpy, fastcluster, and scipy.cluster.hierarchy.fcluster modules (Online Methods, Sec. 6.1). The clustering was performed using the average linkage criterion with a maximum threshold distance imposed. The implementation utilized the Python built-in multi-processing module to parallelize the computations on multiple cores and required approximately 54.4 core-hours and 80GB of memory (computations were performed on a 32 core machine with 250GB RAM) for a dataset containing 100K input antibody sequences. Empirically, the primary bottlenecks for this implementation and dataset were computation time and storage of the distance matrix (note that the embarrassingly parallel nature of this task implies a trivial cost conversion between single and multi-core computations). Full-hierarchical clustering scales comparably in terms of computational complexity; with the threshold imposed, the relative cost is approximately a factor of 50 less than the cost of constructing the distance matrix, however. Although I/O was included in the timing estimates, its contribution in this case was negligible.

Extrapolating the computation to a typical antibody profiling sample containing one million input sequences, the required computational time and storage is expected to reach around 5440 core-hours and 8TB, respectively. Given its poor scalability and efficiency limitations, this implementation is inadequate for large-scale profiling, which for a small clinical vaccine evaluation may consist of dozens of subjects with several longitudinal samples per subject. The goal of the challenge was to optimize and improve the algorithm and its implementation, so that routine data analysis of a large-scale antibody profiling study would become feasible given modest computational resources.

#### 2.1.3 Problem Abstraction and Available Data Sets

A common optimization practice is to convert all or just the computationally-intensive parts of Python codes to languages such as C/C++. We implemented a C++ version of our algorithm (directly translated) and used it in our post-challenge benchmarking analysis (Online Methods, Sec. 6.1). Although the memory footprint for this implementation was comparable to that of the Python implementation, it yielded approximately an order of magnitude reduction in computation time. This computational cost reduction was primarily attributed to the construction of the distance matrix, whereas the computational cost of the clustering itself was comparable in both implementations (this is not surprising, given the underlying implementation of the scipy.cluster.hierarchy.fcluster is written in C). All contest submissions were evaluated relative to the Python (A1) and C++ (A2) benchmarks described above, and the code for both benchmark implementations was provided to the contestants.

We generated datasets of 1K, 5K, 10K and 100K input sequences sampled from true sequences derived from a healthy adult subject and computed their corresponding clustering outputs using the A1 baseline. The output results for these datasets served as the “gold standards” for evaluating the accuracy of the solutions produced in the competition (Online Methods, Sec. 6.1). The training datasets comprised one 1K and one 100K input sequences, the validation datasets comprised two 5K and three 10K input sequences, whereas the testing datasets comprised four 10K and six 100K input sequences. For the given threshold distance, the maximum cluster size for each dataset ranged from 1.6% to 4% of the total number of antibody sequences in the set. The most probable cluster size for each set was unity, however.

#### 2.1.4 Competition Outcome

The competition lasted for 10 days, involved 34 participants, and averaged 6.91 submissions per participant. All contest submissions were evaluated on a 32 core server with 64GB memory, although the top four solutions utilized only a single core. For the given dataset, a majority of participants had submitted solutions that were significantly more computationally efficient than the A2 benchmark, with the winning solutions being orders of magnitude more efficient.

Fig. 1 illustrates the computational cost performance of the A1 and A2 benchmarks, along with the performance envelope for the top four performing solutions as a function of the number of input antibody sequences for *N* up to 1M input sequences. The winning solution was able to perform clustering of 100K sequences in approximately 1.8s on a single core implying a total computational cost reduction (speedup) of 108,800 when compared to the A1 benchmark. Interestingly, the primary bottleneck for this implementation was no longer clustering but rather file I/O. Neglecting file I/O, which accounted for 85% of the computation time, the effective speedup achieved for clustering over the A1 benchmark was approximately 777,000 for 100K sequences^2^. For N up to 1M input sequences, the four solutions required less than approximately 17GB memory, with the winning solution requiring only 0.7GB memory.

**Figure 1:**
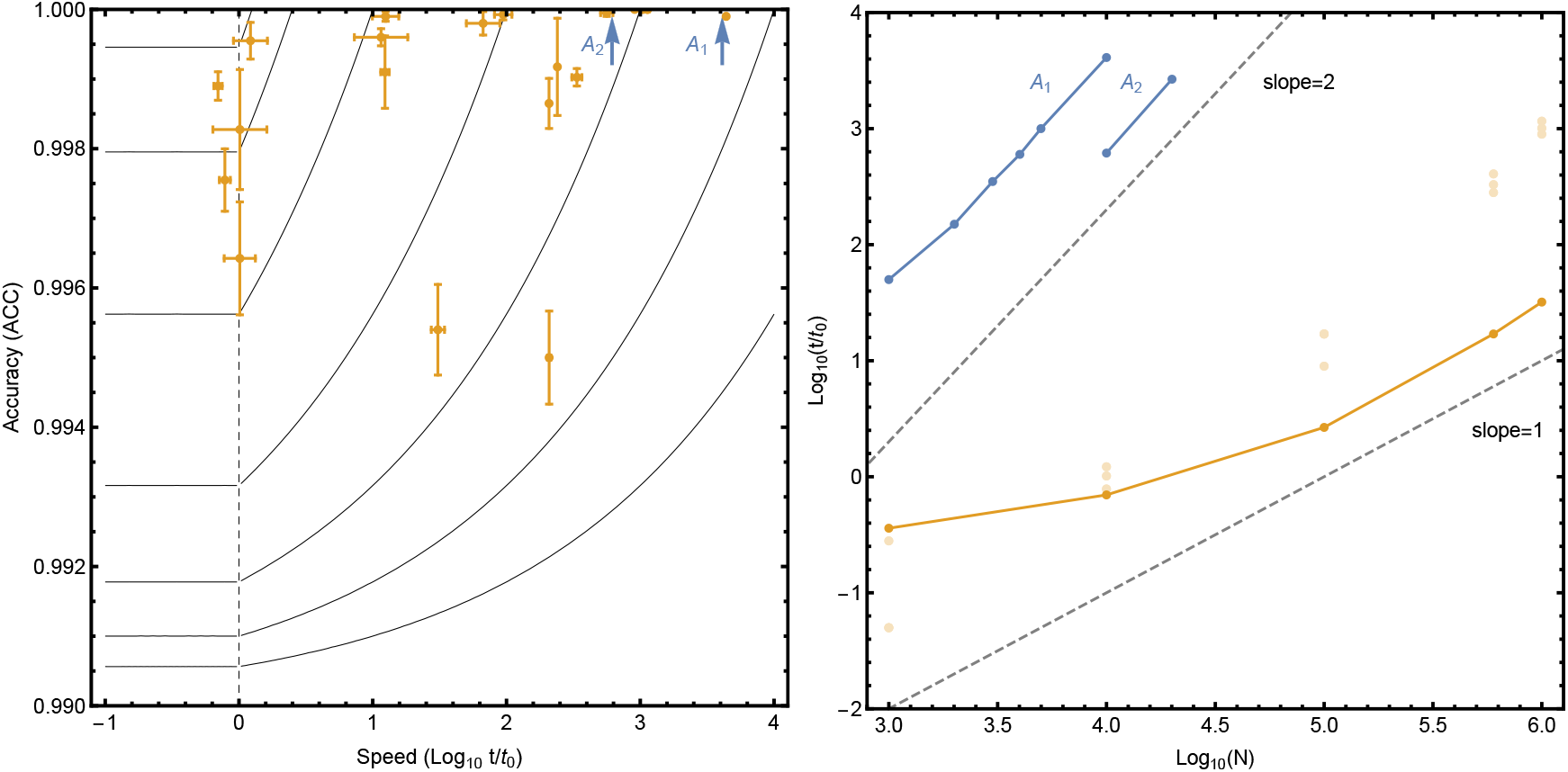
Antibody Challenge outcomes: (Left) Evaluation metric used for Antibody Clustering as a function of time (*t* measured in units of *t*_0_ = 100ms) and accuracy *(ACC),* as determined during the competition. Solutions lying on the isocurves are considered equivalent in quality, as defined by the evaluation metric (see Sec. 6.1). Data points correspond to average performance based on up to four 10K test sets (failed test cases excluded). Also shown is the benchmark algorithm implemented in Python (A1) and C++ (A2); note that benchmark algorithms A1 and A2 have perfect accuracy *(ACC* equal to unity). (Right) Effective computational cost (excluding I/O) on a single core as a function of test set size *N* for Antibody Clustering *(t* measured in units of *to* = 100ms). Dashed lines indicate linear and quadratic scaling behaviors. Results are shown for the benchmark algorithms (A1 and A2) and the top four performing solutions (top solution indicated by solid line and darker data points). The benchmark algorithms exhibit quadratic scaling behavior with test set size, whereas the top algorithm exhibits better than quadratic scaling behavior within the regime considered. The best performance over the entire range of *N* can be achieved using an ensemble of solutions selected based upon whether the test set size is either below or above the crossover value at approximately *N* ~ 0(10^4^).

The winning solution achieved significant improvements in computational efficiency by abandoning generic implementations of hierarchical clustering, which demand computing the full (or half) the distance matrix as input in favor of a problem-specific strategy. The solution exploited several important properties of the data set to dramatically reduce the number of Levenshtein distances computed by noting that:

1. Some families of antibodies are *never* clustered together implying a partitioning of the dataset.
2. Some antibodies are *always* clustered together, forming subclusters.
3. Coarse “superclusters” can be quickly identified and formed from such subclusters.

Once coarse clusters were formed, a final exact clustering was performed for each of the superclusters. In addition to these innovations, the solution exploited SIMD instructions to speed up the calculation, improved the management of memory, and simplified the parsing of input data to minimize I/O operations.

### 2.2 The Connectivity Map Inference Challenge (Broad Institute)

#### 2.2.1 Motivation and Background

The discovery of functional relationships among small molecules, genes, and diseases is a key challenge in biology with numerous therapeutic applications. The Connectivity Map (CMap) allows researchers to discover relationships between cellular states through the analysis of a large collection of perturbational gene expression signatures [12, 13]. These signatures arise from the systematic perturbation of human cell lines with a wide array of perturbagens, such as small molecules, gene knockdown, and over-expression reagents; together, they constitute a massive dataset that can be “queried” by powerful machine learning algorithms to identify biological relationships. Researchers can then use the outcomes of those queries to form hypotheses to guide experimentation [14].

Despite the recent advancements in high-throughput technology, developing massive gene expression data repositories, such as the Connectivity Map, is still considerably challenging due to the cost of doing systematic large-scale gene expression profiling experiments. These experiments typically involve thousands of genes with complex cellular dynamics and nonlinear relationships, thereby usually demanding large sample size. They also involve perturbations that can be performed with a wide array of molecules, at different doses, and in different cellular contexts, thus producing an enormous space of experimentation.

To address these difficulties, the CMap group at the Broad Institute has developed a novel high-throughput profiling technology, called L1000, that makes the data generation process for The Connectivity Map cheaper and faster [13]. This new technology is based on a *reduced representation of the transcriptome* that combines direct measurement of a subset of approximately 1000 genes, called landmarks, with an imputation approach to obtain the gene expression levels for the remaining (non-landmark) genes.

The resulting combination of imputed and directly-measured genes enables the L1000 platform to report on approximately 12,000 genes with accuracy comparable to traditional profiling technologies (e.g., RNA-Seq) but at a fraction of the cost [13]. This improvement has in a few years enabled the Connectivity Map to grow from its initial 500 (Affymetrix) profiles to over one million (L1000) profiles.

While L1000 is a powerful and cost-effective tool, the comparability with externally-generated gene sets remains an open issue. The practice of directly measuring only 1,000 genes (a small fraction of the entire transcriptome) reduces the fidelity of comparisons with externally-generated gene sets, whose overlap with the 1,000 landmarks may be minimal. Improving the imputation accuracy for non-landmark genes is, thus, expected to facilitate comparisons between L1000 and external gene expression data and gene sets, leading to higher-confidence hypotheses.

In addition, while the connectivity analysis of CMap signatures can be restricted to the landmarks only, sometimes this restriction is unfeasible (queries containing no landmarks) or undesirable (leading to poor outcomes, [13]). Improving the imputation accuracy is, therefore, expected to bear additional benefits downstream, increasing the ability of CMap to detect biologically-meaningful connections.

With these goals in mind (improving comparability and the downstream connectivity analysis), the “CMap Inference Challenge” solicited code submissions for imputation algorithms more accurate than the present approach.

#### 2.2.2 Problem Abstraction and Available Data Sets

The following describes the basic imputation approach currently in use by the Connectivity Map:

- Learn the structural correlations between non-landmark and landmark genes using Multiple Linear Regression (MLR) on a reference dataset D with complete data on both landmarks and non-landmarks;
- Perform out-of-sample predictions of the expression values of the non-landmarks in a different target dataset T (i.e., apply the regression function as estimated from D to compute the fitted values of the transcriptional levels for the non-landmark genes in T), using these predictions for imputation.

It has been shown that a MLR approach using L1000 data for imputation achieves considerable accuracy when compared to RNA-Sequencing (RNA-Seq) on the same samples [13]. Nevertheless, alternative imputation approaches could yield performance improvements (e.g., by exploiting possible non-linearities in the data or a larger training dataset).

Contest participants were tasked to explore alternative approaches using an eight-times-larger dataset for their inferential models (Online Methods; Sec. 6.2) with the goal to predict 11,350 non-landmarks from 970 L1000 landmarks. Their models were then evaluated using as ground truth RNA-Seq data profiled on the same samples.

#### 2.2.3 Competition Outcome

Eighty-eight competitors submitted their predictions for evaluation; fifty-five (62%) achieved an average performance higher than the MLR benchmark, with improvements as high as 50%, and with little difference between provisional and system evaluations (Online Methods, Fig. 5), indicating no overfitting.

For the top five submissions, performance improvements were in *absolute* and *relative* accuracy (Fig. 2A-D). The median gene-level rank correlation between the predicted values and the ground truth was 70% higher than the benchmark. The gene-level relative accuracy (i.e., the rank of self-correlation compared to correlations with other genes) was distributed with a lower dispersion (an interquartile range 40-90% lower depending on the submission), resulting in a higher *recall*.^3^

**Figure 2:**
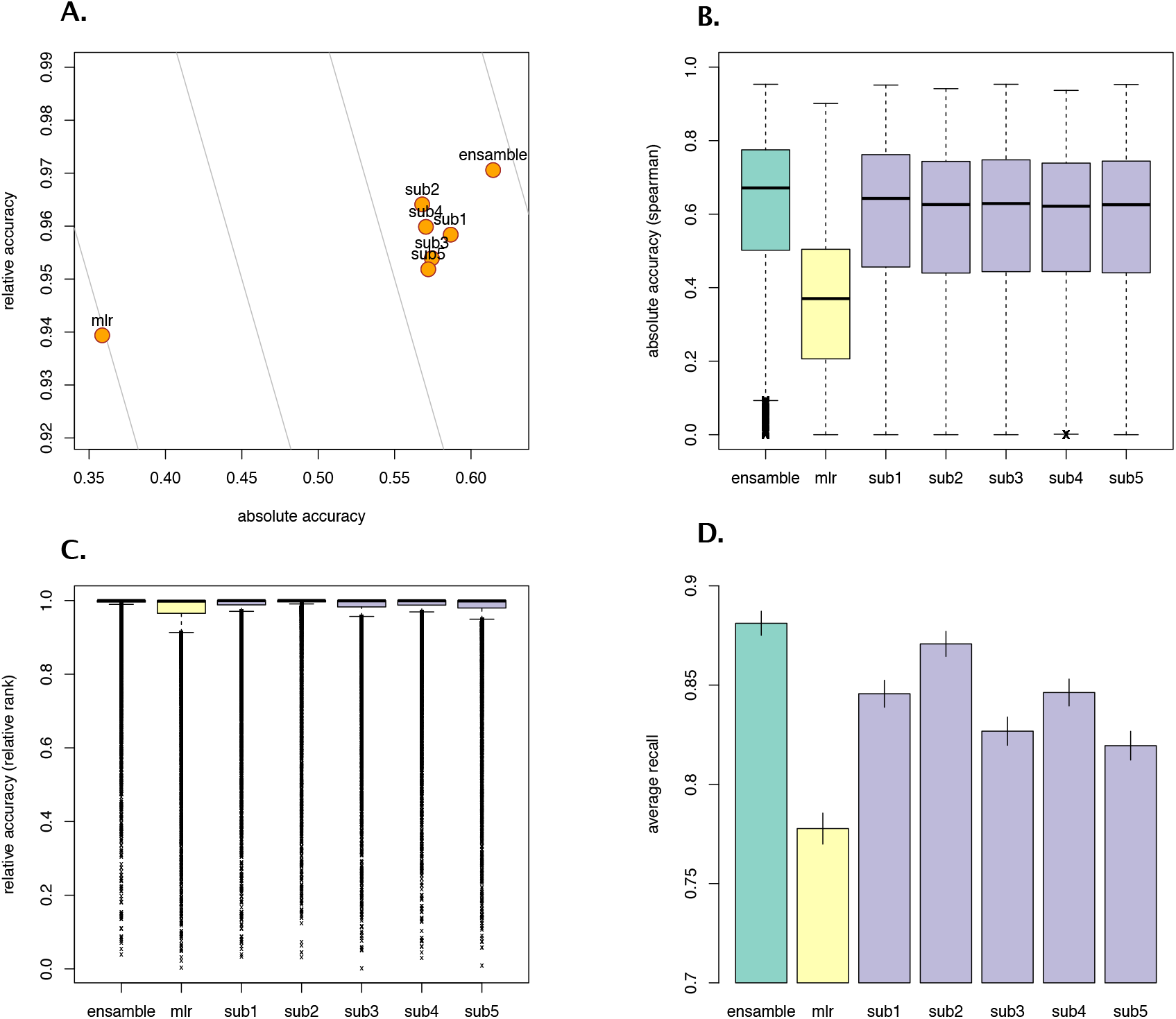
CMap Inference Challenge outcomes: (A) Scatter plot showing the average performance of the top 5 submissions, the benchmark, and an ensemble that takes the best-predicting algorithm from the top 5, where the performance is represented in two dimensions: absolute accuracy (gene-level Spearman correlation) and relative accuracy (gene-level relative rank); curves (light gray) represent locus of points on the plane giving the same aggregate score (curves on the right of this plot represent higher aggregate scores); (B) Boxplot showing absolute accuracy at the gene level for all the top 5 submissions, the benchmark, and the ensemble; (C) Boxplot showing relative accuracy at the gene level for all the top 5 submissions, the benchmark, and the ensemble; and (D) Recall (defined as proportion of genes with a relative accuracy below.95) for all the top 5 submissions, the benchmark, and the ensemble.

Inspection of the methods used by the top five submissions reveals that 4 out of 5 of the top-performing submissions converged to a *K-Nearest Neighbors* (KNN) regression, and only one used a *Neural Network approach.* KNN regression is a non-parametric regression approach that makes predictions for each case by aggregating the responses of its K most similar training examples. Compared to MLR, KNN regression makes weaker assumptions about the shape of the “true” regression function thereby allowing a more flexible representation of the relationship between landmark and non-landmark genes.

Compared to a standard KNN-based imputation model, such as [15], the winning submission presents a few innovations. First, it combines multiple predictions obtained by iteratively applying the same regression function to different data normalization, such as scale, quantile, rank, and ComBat batch normalization [16]. Second, within each iteration, it identifies and combines multiple sets of K nearest neighbors using different similarity measures. These modifications present some potential advantages over a standard KNN method. One potential advantage is to alleviate batch effects thereby controlling for a source of non-biological variation in the data. Another advantage is the identification of the training samples that are most biologically relevant for predicting a given test sample (e.g., same tissue). The net outcome is that the winning approach achieves ¿5% improvement in absolute accuracy and ¿2.6% improvement in the recall over the other top KNN approaches.

We further examined key differences between the winning method and the MLR benchmark by comparing performance on large (100,000) and small (12,000) sample size training datasets. We found that the winning KNN approach outperforms the MLR, although the performance is sensitive to the sample size. When trained on the smaller dataset, the overall performance of the winning KNN approach was higher than the benchmark (10% higher absolute accuracy) but the recall was (5%) lower.

Clustering analysis of the gene-level scores (Fig. 2-E) suggested potential complementarities among the top submissions. We tested this hypothesis by combining the predictions of the top five submissions into an ensemble approach. We used the training dataset to select automatically the best predicting method for each particular gene (the one with the highest combined score). By doing so, we found a strong complementarity between the winning KNN and the Neural Network approach, which were equally selected for over two-thirds of the genes (Fig. 2-F). We then evaluated the resulting predictions on the test set, showing a 2% improved performance over the top performing submission (Fig. 2).

We then examined to what extent a better inference model affects the ability to recover expected connections using CMap data [see 13, for a formal definition of connections]. To test this hypothesis, the winning approach was used to impute non-landmark genes on a dataset of L1000 profiles from about 46,000 samples of multiple perturbagens (Online Methods, Sec. 6.2). We then processed these KNN-imputed data through the standard CMap pre-processing pipeline [13] and queried the resulting signatures, and their MLR equivalents, with a collection of annotated pathway gene sets. Based on the literature, each gene set was expected to connect to at least one of the perturbagens. We compared the distribution of the connectivity scores and corresponding ranks generated using the predictions made by the KNN approach and those made by the benchmark MLR approach. Results show no significant difference in the distributions of these connectivity measures (according to Kolmogorov-Smirnov test).

In conclusion, the top submission achieved substantial improvements in performance over the MLR benchmark, thus it succeeded in achieving a better comparability of CMap inferred data with external data. To facilitate the use of the winning submission for this purpose, the winning code was deployed in the “R” package (“cmapR’) and is currently available at github.com/cmap. At the same time, we found no evidence that the better inference would translate into a more accurate downstream connectivity analysis (higher ability to recover the expected connections), contrary to the initial hypothesis. However, limitations in the contest configuration (scores were not directly based on connectivity) preclude a conclusive statement at the moment, and further investigation is ongoing.

### 2.3 CMap Query Speedup Challenge (Broad Institute)

#### 2.3.1 Motivation and Objective

Researchers often use the Connectivity Map to compare specific patterns of up- and down-regulated genes for similarity to expression signatures of multiple perturbagens (e.g., compounds and genetic reagents) in order to develop functional hypotheses about cell states [13]. These hypotheses can be used to inform areas of research ranging from elucidating gene function to identifying the protein target of a small molecule to nominating candidate therapies for disease.

Given the high potential of this approach, many algorithms that assess transcriptional signatures for similarity have been developed over the years. These algorithms are generally computationally expensive, which may limit their use to relatively small-sized data. The principal problem is that computationally efficient methods, such as cosine similarity, may be inadequate for interpreting gene expression data in general. Moreover, more powerful methods, such as the Gene Set Enrichment Analysis [17], need to perform computationally-expensive tasks, such as ranking genes in each signature by their expression levels to compare gene rank positions individually across signatures. Given the Connectivity Map has recently expanded to over one million profiles [13], this limitation is particularly problematic.

To address this problem, the CMap group developed a fast query-processing algorithm, called Sig-Query. This tool was implemented in MATLAB incorporating a range of optimization techniques to speed up queries on the Connectivity Map, and was available on the online portal CLUE.IO. Overall, the algorithm achieves a good level of performance (in a preliminary analysis, it took about 120 minutes to process 1000 queries with gene sets of size 100 against a signature matrix of 470,000 signatures). Even so, execution time and memory requirements are still a potential barrier to adoption for the Connectivity Map.

To further the development of query algorithms for CMap data, the “CMap Query Speedup Challenge” solicited code submissions for fast implementations of the present CMap query methodology.

#### 2.3.2 Problem Abstraction and Available Data Sets

Following the query methodology described in Sec. 6, one bottleneck lies at rank-ordering all signature values and subsequently walking down the entire list of G genes in a signature to compute the running sum statistics over all possible pairs of S signatures and queries. Using the quick-sort algorithm, this task has a computational complexity of *O*(*S × G* log(*G*)) in the average case. To save time, however, results can be stored on disk, as in the current implementation, with adjustments to try to minimize the cost of access to disk memory. The other very burdensome task involves computing the running sums for all genes in each signature, which has a computational complexity upper bound that scales as *O*(*S × G*) per query.

For this contest, participants had to address these problems and the performance of their methods was evaluated on 1000 queries to be run on the whole CMap signature matrix, which has expression values for > 470k signatures and > 10k genes (Online Methods, Sec. 6.3).

#### 2.3.3 Competition Outcome

The competition resulted in 33 participants making 168 code submissions. All final submissions were evaluated on the holdout query dataset on a server with 16 cores. Results showed significant speed improvements over the benchmark: the median speedup was of 23 × with at least two submissions achieving speedups beyond 60 ×.

Comparison of performance between 16-core and single-core evaluations showed that *multithreading* alone accounted for a large fraction of the gains in performance over the benchmark (Fig. 4-A): the median speedup difference between single and multi core was 18 ×, accounting for 70% of the final median speedup over the benchmark.

**Figure 3:**
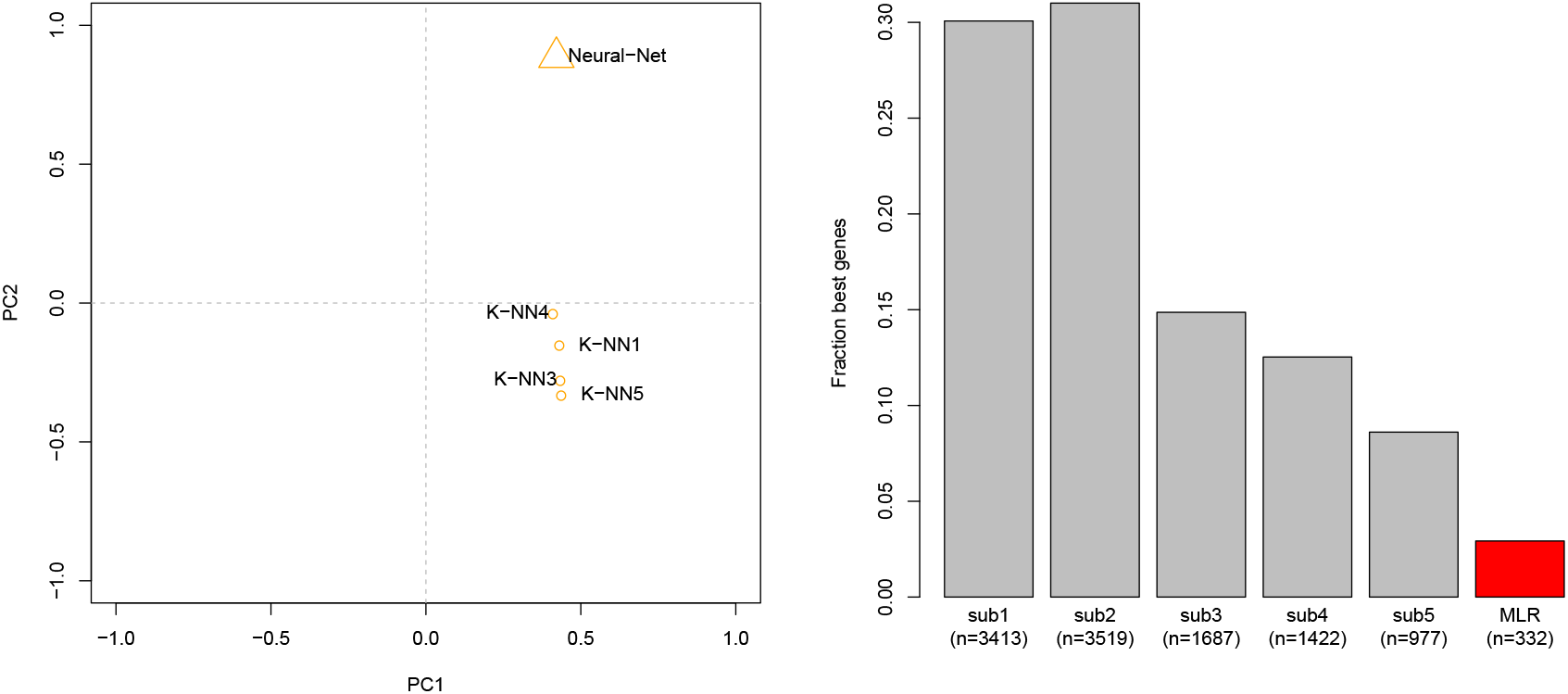
Left: Plot showing results of the Principal Component Analysis (PCA) of the gene-level combined scores of all top five submissions for the CMap Inference Challenge. While all top submissions are clustered together in the first factor of PCA, they differ in the second factor, separating into different “clusters” the Neural Network approach (submission 2) and the other KNN based approaches (submissions 1, 3, 4, and 5). Right: Barplot showing the proportion (and count in parenthesis) of gene predictions used to form the ensemble, which uses the training dataset to select the best predicting algorithm from the MLR benchmark and the top 5 submissions (i.e., the algorithm achieving the highest combined score for a particular gene) for the CMap Inference Challenge.

**Figure 4:**
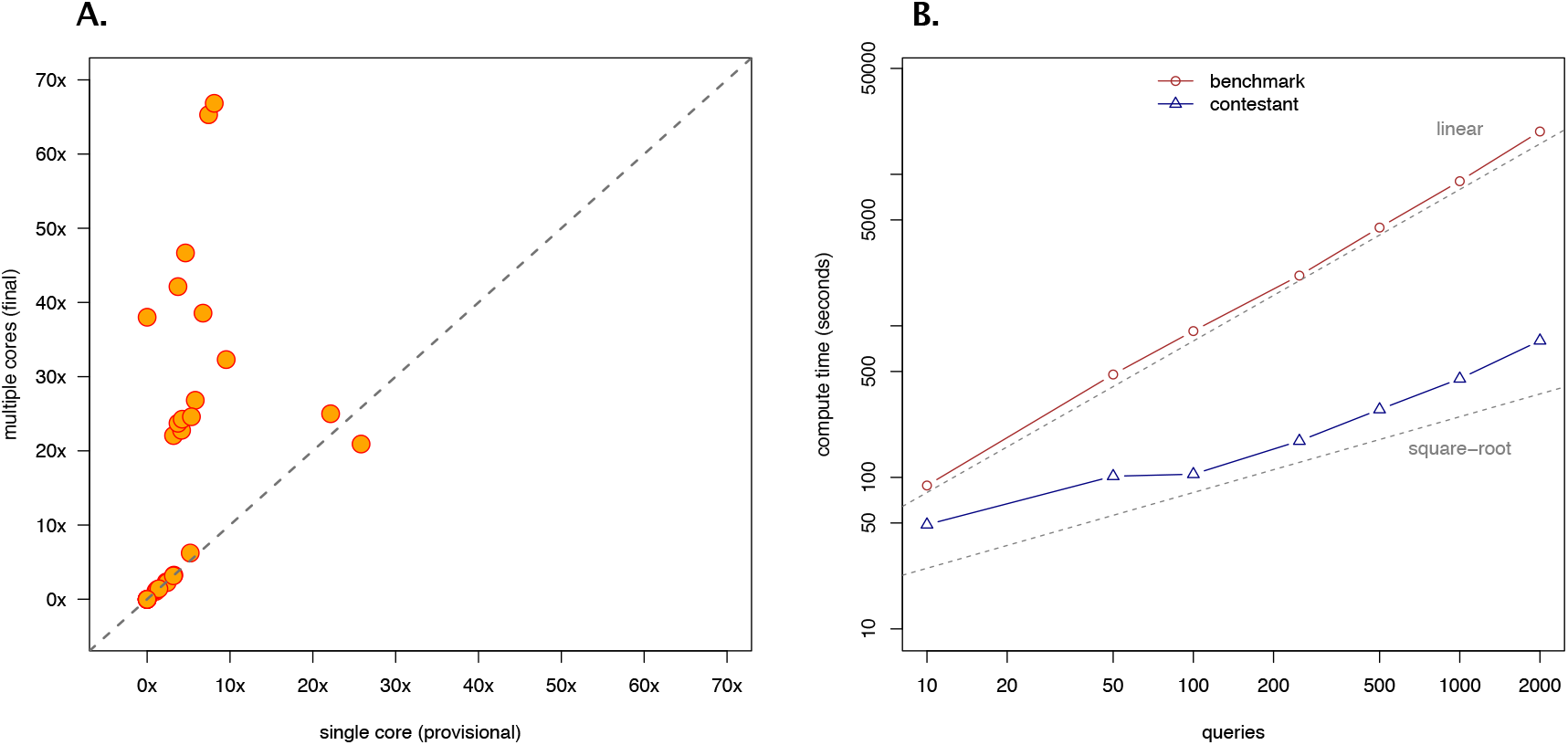
CMap Query Speedup Challenge Results: (A) Scatter plot showing the relationship between speedups computed on the participants’ last submission with one single core (x-axis) and those with multiple cores (y-axis); observations lying on the 45 degrees line correspond to participants who did not use multi-threading in their final submission (with small variation around the line due to differences in the set of queries used to compute the speedups); (B) Plot showing differences in the scaling behavior between the benchmark and the top submission (on a log-log scale); for both, the displayed computation times are a median of 10 replicates.

Beyond multithreading, analysis of the scaling properties of the winning submission showed a computational time complexity that scales as the *square-root* of the number of queries (Figure 4-B), which represents a major improvement compared to the benchmark’s *linear* scaling.

The winning submission also showed substantial performance improvements in the reading time —the time to load in memory the absolute values and rank-ordered positions of genes in the CMap signature matrix: the overall reading time of the winning submission was just below 10 seconds, which represents a 50× speedup compared to the 500 seconds used by the benchmark (Online Methods, Fig 6).

**Figure 5:**
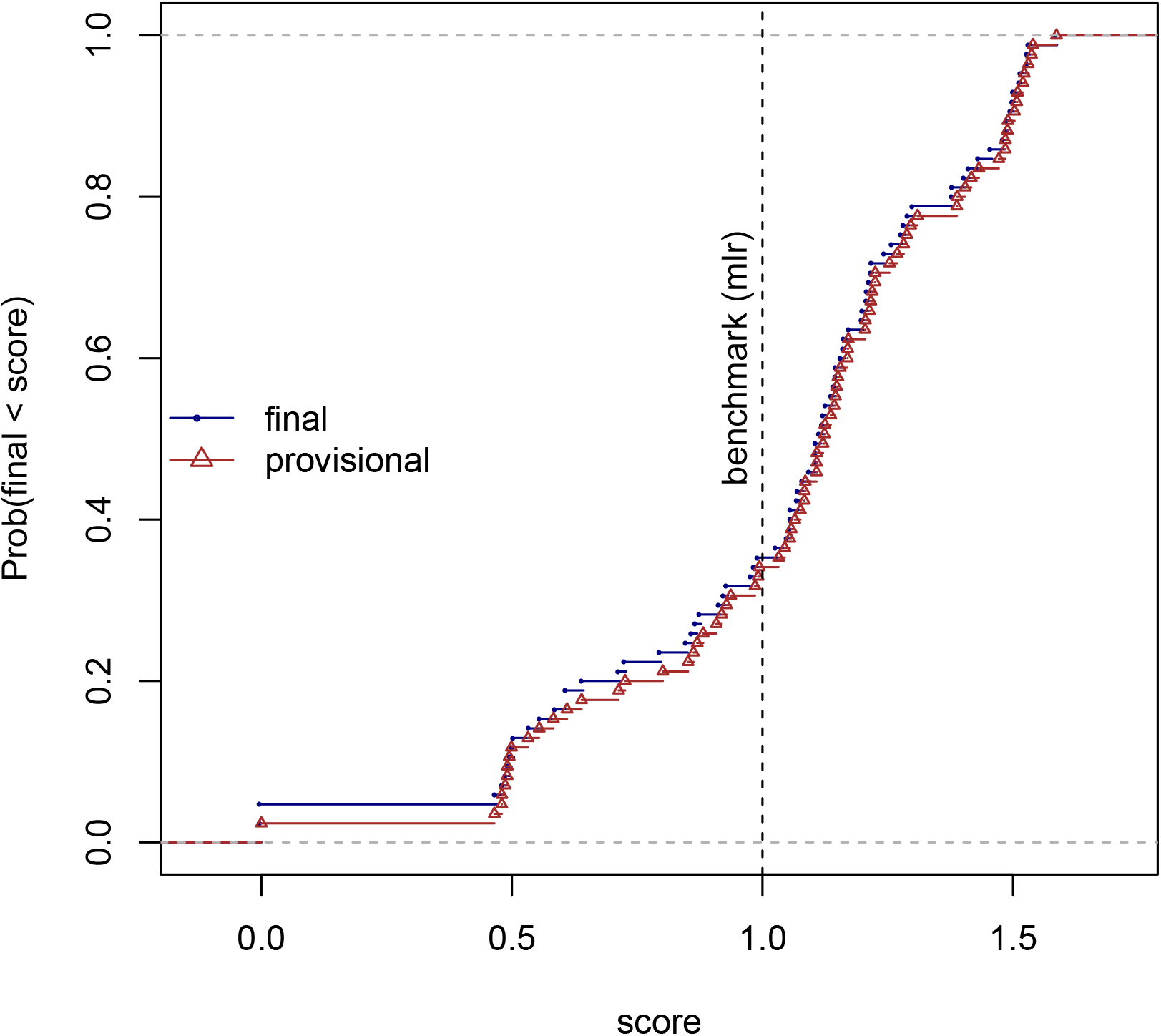
Plot showing the empirical distribution functions for the provisional and final scores computed on all the final submissions for the CMap Inference Challenge. The two distribution curves overlap quite well indicating no overfitting.

**Figure 6:**
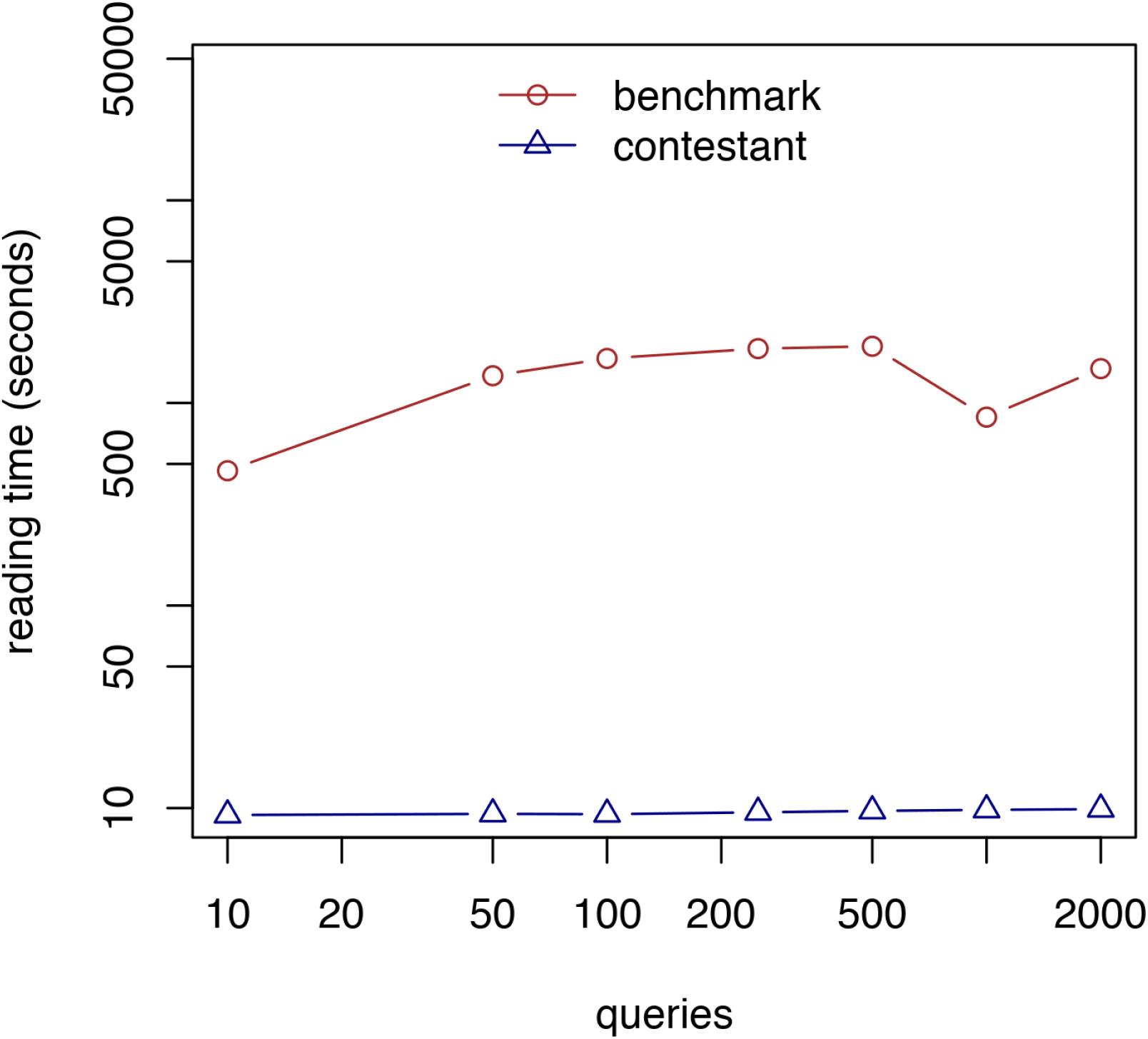
Read time as a function of number of queries on a log-log scale; dots plot median time out of 10 replicates for each dataset for CMap Query Speedup Challenge.

Close examination of the codes of the top submission revealed multiple optimization adjustments that can account for all these improvements. These adjustments often overlap with those of the other submissions, thus making hard any clear-cut categorization. Consequently, we report instead the areas of optimization that we believe are responsible for most of the performance improvement over the benchmark (beyond multithreading and the recourse to low-level programming languages, such as C++, instead of MATLAB). Essentially, these are:

1. Efficient data storage techniques in order to maximize the available cache memory for each thread.
2. Streaming SIMD Extensions (SSE) technology to execute multiple identical operations simultaneously for each thread.

One of the ways by which the winning submission achieved its exceptional performance was by loading the entire signature matrix in the cache memory. Cache memory is indeed the fastest memory in the computer, albeit of very limited capacity. To minimize memory usage, the winning contestant stored the CMap signature matrix at a lower precision than the benchmark (32-bit single precision floating for the scores and 16-bit integers for the ranks), with essentially no loss in accuracy. Precision reduction alone, however, was insufficient to fit level 1 cache memory (the fastest cache memory available) due to the large extent of queries and gene sets to be processed per signature. So, it developed a clever system of matrices to efficiently store the indexes and partial sums for each gene in a query. The resulting algorithm made a much more efficient use of memory compared to the benchmark.

The other major improvement is related to SSE, which is a set of instructions that allows the processor to compute the same operation on multiple data points simultaneously [18]. The winning submission used SSE to form batches of 4 genes and simultaneously compute the rank positions of these genes, thus reducing by approximately a factor of one-fourth the time of each query.

As a result, the winning code submission for the CMap Query Speedup Challenge was deployed in the online portal CLUE.io and is now currently available as an option to users in the Query App (Online Methods, Fig. 7). This improved algorithm has also enabled CLUE to support batch queries, allowing users to execute multiple queries in a single job, all via the CLUE user interface.

**Figure 7:**
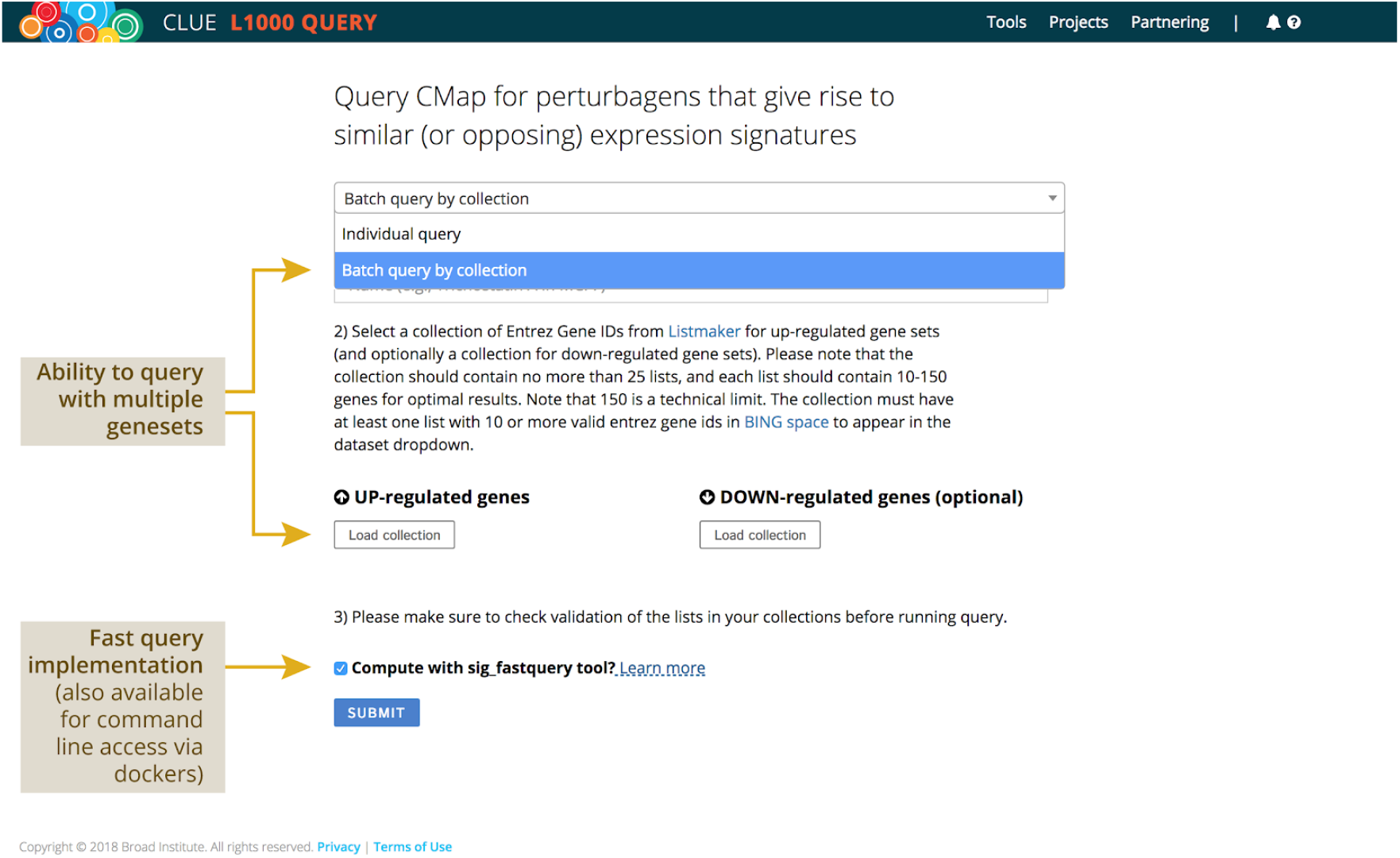
Screenshot of the implementation of the winning code submission for the CMap Query Speedup Challenge in the online portal CLUE.io, where the code is currently available as an option to users in the Query App (“compute with sig_fastquery tool”).

## 3 Discussion

This work demonstrates how researchers in computational biology and bioinformatics have utilized open innovation competitions to make inroads on a variety of computational roadblocks in their work. While participants in these competitions may not possess the domain knowledge to solve every research problem faced by scientists, their unique skills can be leveraged to attack well-defined tasks (e.g., algorithm improvement to maximize computational speed) resolving specific issues or bottlenecks to the research process.

We highlight a few key advantages over traditional approaches. First, competitions enable rapid yet broad exploration of the solution space of the problem. This broad exploration opens the possibility of discovering high-performing solutions, which may lead to breakthroughs in the field. Second, competitions give researchers access to multiple complementary solutions. Thus, they create opportunities to boost performance even further with ensemble techniques whereby different approaches are combined based upon their strengths in different regimes or for different subsets of data.

Our results indicate that these gains arise from the efforts of out-of-the-field participants instead of the community of practitioners and domain experts. Thus, on the one hand, we know open innovation competitions are a great tool to solicit community-based effort (e.g., Dream Challenges); on the other, we show that the potential for their use in biology and other life sciences goes beyond the size and availability of the community of researchers connected to the problem.

By defining appropriate objective functions to guide the competitors’ efforts, researchers have indeed the flexibility to pursue vastly different problems or even decide to tackle multiple aspects of the same problem concurrently via a composite objective function. This kind of problem definition, however, can be difficult. It requires knowledge of the aspects that are critical to the problem and for which improvements are quantifiable and achievable, understanding of ways to trade-off improvements in one dimension for another, and the ability to abstract the original problem from its domain to attract broad participation (knowing that domain-specific information hurts participation but also offers competitors insights on how to solve the problem).

We have shown some of the challenges encountered in addressing these issues for two very general types of computational problems: “code development” and “machine learning.”

Code development problems typically boil down to improving existing computational algorithms that have well-defined inputs and outputs under a variety of constraints (e.g., speeding up the computation, without exceeding memory limits). Here, although performance improvements are quantifiable and can be checked by test cases, other relevant aspects (e.g., robustness) are less so. This restriction forces researchers to take additional steps after the contest to validate methodologies and integrity of solutions beyond the limited test cases considered during the competition. These steps typically include ensuring security (understanding what the code does and ensuring it is not malicious) and legality of the produced codes (that they are original or properly licensed) before integration and deployment.

Machine learning problems focus on more exploratory questions, such as producing predictive models that describe an existing data set, yet are generalizable to new data. These problems are typically multi-dimensional, given the wide range of potential applications, and are sometimes hard to quantify (e.g., measurements that offer only a partial picture of biological states to model). As a result, competitions addressing these problems typically involve interpretation and further evaluation of methodologies, exploration of possible complementarities, and understanding strengths and pitfalls of solutions in comparison to known methodologies along dimensions not considered within the competition.

Our study suggests that although competitions offer the flexibility to address both kinds of problems, the associated post-competition efforts can be quite different. In both cases, handling these additional tasks requires expertise that itself may not be available to the end-user, although they may be addressed in part by subsequent competitions; thus, keeping a modular design strategy appears beneficial. For ML problems, further efforts (comparable to those devoted to replicating the methods used in another study) are often needed to evaluate and assimilate the new knowledge produced. Further research to examine the outer limits of scientific problem solving through contests is necessary.

We conclude by mentioning a few additional empirical contexts in which open innovation competitions seem promising. One is the assignment of functional attributes of small molecules and genes, such as predicting the mechanism of action of a compound. While many algorithms have been independently developed for this purpose, a broad exploration is often out of reach to individual laboratories. On the development side, research in biology is often impeded by bottlenecks in the memory storage and transmission of genetic data, which are critical and quantifiable, thus an open innovation competition seems an effective way to overcome these bottlenecks.

## 4 Acknowledgments

This work was funded by the Eric and Wendy Schmidt Foundation, NASA Center of Collaborative Excellence, and the Kraft Precision Medicine Accelerator & Division of Research and Faculty Development at the Harvard Business School. The Scripps competition was funded by N.I.H. Grant Number 5UM1 AI100663. The CMap competitions were supported in part by the NIH Common Funds Library of Integrated Network-based Cellular Signatures (LINCS) program U54HG008699 and NIH Big Data to Knowledge (BD2K) program 5U01HG008699.

## 6 Online Methods

### 6.1 Antibody Clustering Challenge (Scripps)

#### 6.1.1 Competition Evaluation

Evaluation of a multi-task challenge such as this, which aims to yield performance gains in speed and memory use while maintaining a high degree of accuracy, requires some care due to the unavoidable interplay between performance characteristics which is inherent to scalar metrics. Therefore, we developed a weighted scoring function to evaluate the contestants’ solutions against the gold standards. For a given dataset, scores were assigned to each algorithm based on

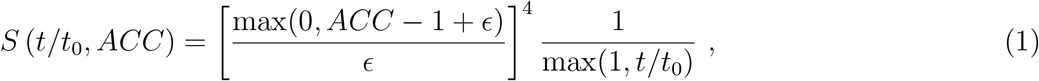

where *t* is the computation time required for a given test set on a 32 core server, *ACC* is the associated accuracy of the computation compared to the gold standards, *ϵ* = 10^−2^, and *t*_0_ = 100ms. Solutions that exceeded a memory use threshold of 3GB received a score of zero. The accuracy (ACC) was determined from the similarity between two clustering based on an adaptation of the Rand index [19]. The final evaluation was based on the average performance on multiple test sets, after normalizing the scores for each test set by the top-performing solution. Fig. 1 illustrates the average performance of all final submissions as a function of computational cost and accuracy for 10K hold-out datasets, as compared to the A1 and A2 benchmarks.

#### 6.1.2 Data Access

The training, validation and test data sets,as well as the Python and C++ benchmark clustering algorithms are available at https://github.com/SuLab/Antibody-Clustering-Challenge.

### 6.2 CMap Inference Challenge (Broad Institute)

#### 6.2.1 Competition Evaluation

To assess imputation accuracy, we created an evaluation metric that balanced both absolute and relative measures of accuracy. Let *P_ij_* represent the predicted expression levels for a set of samples, indexed by *i* = 1,…,*N*, and non-landmark gene labels, indexed by *j* = 1,…,*M* (for this study, *M* = 11350). Similarly, let *G_ik_* represent the true (measured) expression levels for the same set of samples and non-landmark genes, indexed *k* = 1,…,. For each pair of gene labels *j,k,* construct the Spearman rank correlation matrix elements *ρ_jk_*(*P,G*) between prediction and truth data across all the samples. For a given gene label *j*, let *R_j_*(*P,G*) be the relative rank of the correlations *ρ_ij_*(*P,G*) where *k* = *j* with respect to the remaining correlations *k* ≠ *j*. The score attributed to each gene-level prediction is given by an equally-weighted average of the Spearman correlation between prediction and truth data, and the relative rank of the correlation when compared with the correlations associated with the remaining genes:

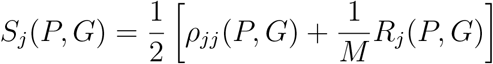

For this competition, gene-level scores were normalized with respect to a benchmark algorithm, which was based on a linear regression model. Let *B_ij_* represent the predictions made by the benchmark algorithm and *S_j_*(*B, G*) represent the gene-level scores associated with the benchmark predictions. The final quality of the predictions was determined based on the average:

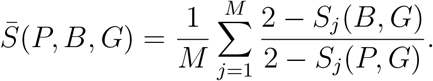

#### 6.2.2 Data Access

All the datasets related to this challenge can be found at the Gene Expression Omnibus (GEO) data repository, www.ncbi.nlm.nih.gov/geo/download/?acc=GSE92743. Test samples were generated in collaboration with the Genotype Tissue Expression (GTEx) project (www.gtexportal.org/home/).

A brief description of the main files is as follows:

- *Training:* Affymetrix data of 12,320 gene expressions for 100,000 samples used by contestants for building their models, ftp://ftp.ncbi.nlm.nih.gov/geo/series/GSE92nnn/GSE92743/suppl/GSE92743_Broad_Affymetrix_training_Level3_Q2NORM_n100000x12320.gctx.gz.
- All of the gene expressions (directly measured & inferred) of L1000 measurements on 3,176 GTEx samples, ftp://ftp.ncbi.nlm.nih.gov/geo/series/GSE92nnn/GSE92743/suppl/GSE92743_Broad_GTEx_L1000_Level3_Q2NORM_n3176x12320.gctx.gz.
- *Validation Set:* Subset of L1000 measurements (directly measured & inferred) on 650 samples randomly selected for provisional scoring, ftp://ftp.ncbi.nlm.nih.gov/geo/series/GSE92nnn/GSE92743/suppl/GSE92743_Broad_GTEx_L1000_Test_Level3_Q2NORM_n650x12320.gctx.gz.
- *Test Set (inputs):* Subset of L1000 measurements (directly measured & inferred) on 1000 samples randomly selected for final scoring, ftp://ftp.ncbi.nlm.nih.gov/geo/series/GSE92nnn/GSE92743/suppl/GSE92743_Broad_GTEx_L1000_Holdout_Level3_Q2NORM_n1000x12320.gctx.gz.
- *Test Set (ground truth):* All of the RNA-seq measurements on 3,176 GTEx samples, which were used as ground truth for provisional and final scoring, ftp://ftp.ncbi.nlm.nih.gov/geo/series/GSE92nnn/GSE92743/suppl/GSE92743_Broad_GTEx_RNAseq_Log2RPKM_q2norm_n3176x12320.gctx.gz.
- Matrix of weights used in current CMap L1000 inference model ftp://ftp.ncbi.nlm.nih.gov/geo/series/GSE92nnn/GSE92743/suppl/GSE92743_Broad_OLS_WEIGHTS_n979x11350.gctx.gz.

The dataset with Affymetrix gene expressions that was used to train originally the CMap L1000 MLR inference model can be downloaded from the GEO data repository, https://www.ncbi.nlm.nih.gov/geo/download/?acc=GSE92742

- File *DS-GEO-nî2031x22268.gctx*in auxiliary dataset (ftp://ftp.ncbi.nlm.nih.gov/geo/series/GSE92nnn/GSE92742/suppl/GSE92742_Broad_LINCS_auxiliary_datasets.tar.gz).

### 6.3 CMap Query Speedup Challenge (Broad Institute)

#### 6.3.1 CMAP Query Methodology

To assess the similarity between a query and a signature, the methodology used by CMap is based on the weighted Kolmogorov-Smirnov enrichment statistic [13]. This statistic reflects the degree to which genes in the query *q* are overrepresented at the top or bottom of the signature *s*.

Let a signature s be a vector of *G* differential expression values *x_g_* for each gene *g* with *g* = 1, 2,…, *G*.

Let a query *q* be a list *q* = {*G*_up_, *G*_down_} of two disjoint gene sets *G*_up_ and *G*_down_ indicating which genes are expected to be up- and down-regulated (the biological state of interest).

The weighted Kolmogorov-Smirnov enrichment statistic is computed as follows:

1. For each gene set Gĳ, compute the sum of values at all positions of s that correspond to genes in *G_j_*:

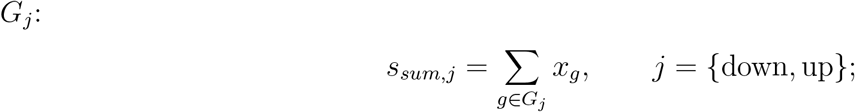
2. Sort *s* in descending order to obtain a rank-ordered signature *s^r^*
3. For each gene set *G_j_*, compute a running sum statistic *rss_g,j_* that walks down the rank-ordered signature *s^r^* and, at each rank position *r_g_* corresponding to a gene *g*, is incremented by a factor *r_g_*/*s_sum,j_* when *g* ∈ *G_j_*, otherwise is reduced by 1/(*G* – |*G_j_*|);
4. Compute the maximum deviation of the running statistic *rss_gmax,j_* from zero for both gene sets *j* = down, up
5. Finally, if the deviations corresponding to each gene set have a different sign, return the average absolute deviation; otherwise return zero (i.e., no similarity is found).

Extending this procedure to a database with *S* signatures and *Q* queries is straightforward and it consists of iteratively applying the above procedure to all pairs of signatures and queries.

#### 6.3.2 Competition Evaluation

For a given set of queries, let *sp* = *t/b* be the speedup over the benchmark, which is defined by the ratio between *t* the time (in seconds) a participant’s submission takes to complete all the queries and *b* the runtime of the existing SigQuery tool for the same task. All code submissions were timed on the same server and rank-ordered based on the score

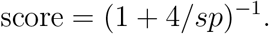

The choice of using the *runtime* for scoring, instead of the *process time* (e.g., CPU time), deserves further comment. If the scoring function was based on the process (or CPU) time, participants would have had an incentive not to use *multithreading* techniques in their submissions, given the additional inter-processor communication overheads. By contrast, a scoring function based on runtime speedups would encourage competitors to use *multithreading,* which was available to everyone in the final evaluation process (i.e., all codes were evaluated on the same machine with 16 cores). So, the choice of using runtime for scoring was essentially to encourage implementations of multithreaded solutions.

In addition to speed, submissions had to be considered sufficiently accurate to be eligible for prizes. We measured accuracy as the lowest absolute deviation in the Kolmogorov-Smirnov statistics between those obtained by the competitor and those computed by the SigQuery tool. Submissions were thus required to have their lowest absolute deviation below a given threshold (i.e., 0.0001). Note that we focused on the lowest absolute deviation of all the Kolmogorov-Smirnov statistics that were computed on each set of up- and down-regulated genes (i.e., before the possible normalization to zero, as described above) separately. This choice reflects considerations about the possibility that strategies for optimizing data storage may result in small losses in precision that may, in turn, cause random changes in the sign of the Kolmogorov-Smirnov statistics, when the absolute value of the statistic is low. An additional requirement was, hence, to allow a maximum of 1000 such differences.

#### 6.3.3 Data Access

The gene sets used for the queries in this challenge were obtained by downloading public Affymetrix data from the National Center for Biotechnology Information’s GEO repository and performing comparative marker selection between case and control samples in order to identify differentially expressed genes [20]. For the convenience of analysis, we fixed the total size of the gene sets to 100, a number which was intended to mimic the typical size of queries.

As with the other challenges, the dataset of queries was randomly split into training, validation, and test sets of size 250, 250, and 500. Queries in the training and validation sets were provided as CSV files, each containing gene set identifiers (rows) and a list of identifiers of individual genes contained in the gene set (columns). The test dataset was withheld and used to validate the final submissions.

Competitors had access to the whole CMap signature matrix that was stored and distributed as a series of CSV files containing the matrix of differential gene expression values and the rank-ordered matrix that was pre-computed by sorting the signature matrix in descending order.

## Footnotes

1 Although multiple motivations, such as the enjoyment for the competition itself, may be at play as well.

2 Further improvement of the winning algorithm was achieved subsequent to the competition, in part by aggressively optimizing the file I/O. The improvements resulted in another 30-fold reduction in the total computational cost of the algorithm (including I/O) on the 100K dataset.

3 All the above differences were highly statistically significant (*p* < 0.001), according to a pairwise paired Wilcoxon Signed Rank Test with Bonferroni correction.

## References

[1] Alex J Hughes, Joseph D Mornin, Sujoy K Biswas, Lauren E Beck, David P Bauer, Arjun Raj, Simone Bianco, and Zev J Gartner. Quanti.us: a tool for rapid, flexible, crowd-based annotation of images. Nature methods, 15(8):587–590, July 2018.

[2] Seth Cooper, Firas Khatib, Adrien Treuille, Janos Barbero, Jeehyung Lee, Michael Beenen, Andrew Leaver-Fay, David Baker, Zoran Popovic, and Foldit players. Predicting protein structures with a multiplayer online game. Nature, 466(7307):756, August 2010.

[3] Karim R Lakhani, Kevin J Boudreau, Po-Ru Loh, Lars Backstrom, Carliss Baldwin, Eric Lonstein, Mike Lydon, Alan MacCormack, Ramy A Arnaout, and Eva C Guinan. Prize-based contests can provide solutions to computational biology problems. 31(2):108–111, 2013.

[4] Steven M Hill et al. Inferring causal molecular networks: empirical assessment through a community-based effort. Nature methods, 13(4):310–318, April 2016.

[5] Julio Saez-Rodriguez, James C Costello, Stephen H Friend, Michael R Kellen, Lara Mangravite, Pablo Meyer, Thea Norman, and Gustavo Stolovitzky. Crowdsourcing biomedical research: leveraging communities as innovation engines. Nature reviews. Genetics, 17(8):470–486, July 2016.

[6] James C Costello et al. A community effort to assess and improve drug sensitivity prediction algorithms. Nature Biotechnology, 32(12):1202–1212, December 2014.

[7] Lars Bo Jeppesen and Karim R Lakhani. Marginality and problem-solving effectiveness in broadcast search. Organization science, 21(5):1016–1033, 2010.

[8] Raymond H Mak, Michael G Endres, Jin H Paik, Rinat A Sergeev, Hugo Aerts, Christopher L Williams, Karim R Lakhani, and Eva C Guinan. Use of crowd innovation to develop an artificial intelligence-based solution for radiation therapy targeting. Submitted to JAMA Oncology.

[9] William D. Lees and Adrian J. Shepherd. Studying Antibody Repertoires with Next-Generation Sequencing, pages 257–270. Springer New York, New York, NY, 2017.

[10] Pascale Mathonet and Christopher Ullman. The application of next generation sequencing to the understanding of antibody repertoires. Frontiers in Immunology, 4:265, 2013.

[11] D. R. Burton and J. R. Mascola. Antibody responses to envelope glycoproteins in HIV-1 infection. Nat. Immunol., 16(6):571–576, Jun 2015.

[12] Justin Lamb et al. The Connectivity Map: Using Gene-Expression Signatures to Connect Small Molecules, Genes, and Disease. Science, 313(5795):1929–1935, September 2006.

[13] Aravind Subramanian et al. A Next Generation Connectivity Map: L1000 Platform and the First 1,000,000 Profiles. Cell, 171(6):1437–1452.e17, April 2018.

[14] Xiaoyan A Qu and Deepak K Rajpal. Applications of Connectivity Map in drug discovery and development. Drug Discovery Today, 17(23–24):1289–1298, December 2012.

[15] Trevor Hastie, Robert Tibshirani, Gavin Sherlock, Michael Eisen, Patrick Brown, and David Bot-stein. Imputing Missing Data for Gene Expression Arrays. 1999.

[16] Jeffrey T Leek, W Evan Johnson, Hilary S Parker, Andrew E Jaffe, and John D Storey. The sva package for removing batch effects and other unwanted variation in high-throughput experiments. Bioinformatics (Oxford, England), 28(6):882–883, March 2012.

[17] Aravind Subramanian, Pablo Tamayo, Vamsi K Mootha, Sayan Mukherjee, Benjamin L Ebert, Michael A Gillette, Amanda Paulovich, Scott L Pomeroy, Todd R Golub, Eric S Lander, and Jill P Mesirov. Gene set enrichment analysis: a knowledge-based approach for interpreting genome-wide expression profiles. Proceedings of the National Academy of Sciences, 102(43):15545–15550, October 2005.

[18] S K Raman, V Pentkovski, and J Keshava. Implementing streaming SIMD extensions on the Pentium III processor. IEEE Micro, 20(4):47–57, 2000.

[19] William M. Rand. Objective criteria for the evaluation of clustering methods. Journal of the American Statistical Association, 66(336):846–850, 1971.

[20] Joshua Gould, Gad Getz, Stefano Monti, Michael Reich, and Jill P. Mesirov. Comparative gene marker selection suite. Bioinformatics, 22(15):1924–1925, 2006.

